# Functional RNA Structures in the 3’UTR of Tick-Borne, Insect-Specific and No-Known-Vector Flaviviruses

**DOI:** 10.1101/565580

**Authors:** Roman Ochsenreiter, Ivo L. Hofacker, Michael T. Wolfinger

## Abstract

Untranslated regions (UTRs) of flaviviruses contain a large number of RNA structural elements involved in mediating the viral life cycle, including cyclisation, replication, and encapsidation. Here we report on a comparative genomics approach to characterize evolutionarily conserved RNAs in the 3’UTR of tick-borne, insect-specific and no-known-vector flaviviruses *in silico*. Our data support the wide distribution of previously experimentally characterized exoribonuclease resistant RNAs (xrRNAs) within tick-borne and no-known-vector flaviviruses and provide evidence for the existence of a cascade of duplicated RNA structures within insect-specific flaviviruses. On a broader scale, our findings indicate that viral 3’UTRs represent a flexible scaffold for evolution to come up with novel xrRNAs.

## 1. Introduction

Flaviviruses are small, single-stranded positive-sense RNA viruses that are typically transmitted between arthropod vectors and vertrebrate hosts. They are endemic in tropic and sub-tropic regions and represent a global health threat, although humans are considered dead end hosts in many cases.

The genus *Flavivirus* within the *Flaviviridae* family comprises more than 70 species, which are organized into four groups, each with a specific host association: Mosquito-borne flaviviruses (MBFVs) and tick-borne flaviviruses (TBFVs) spread between vertebrate (mammals and birds) and invertebrate (mosquitoes and ticks) hosts, whereas insect-specific flaviviruses (ISFVs) replicate specifically in mosquitoes and no-known-vector flaviviruses (NKVs) have only been found in rodents and bats, respectively. This natural host-range-based classification is in good agreement with sequence-based phylogenetic clustering, mainly because all flaviviruses share a common genome organization [1]. Conversely, epidemiology, disease association [2] and transmission cycles [3] are fundamentally different among different flavivirus groups.

Emerging and re-emerging MBFVs such as Dengue virus (DENV), Japanese encephalitis virus (JEV), West Nile virus (WNV), Yellow fever virus (YFV) or Zika virus (ZIKV) are the causative agents of large-scale outbreaks that result in millions of human and veterinary infections every year [4]. Likewise, tick-borne encephalitis virus (TBEV), Powassan virus (POWV) and other members of the tick-borne serocomplex are neuropathogenic agents that cause a large number of infections every year, resulting in a massive incidence increase since the 1970ies [5]. Consequently, much research effort has gone into studying MBFV and TBFV biology, biochemistry and phylogeny [6]. The two remaining groups, ISFVs and NKVs, however, have received limited attention in the research community, mainly because they are generally not associated with human or veterinary disease and therefore are still underrepresented in the literature. The phylogenetic relationship among the four ecological flavivirus groups is shown in Fig 1. Table A1 lists all viral species considered in the present study.

**Figure 1.**
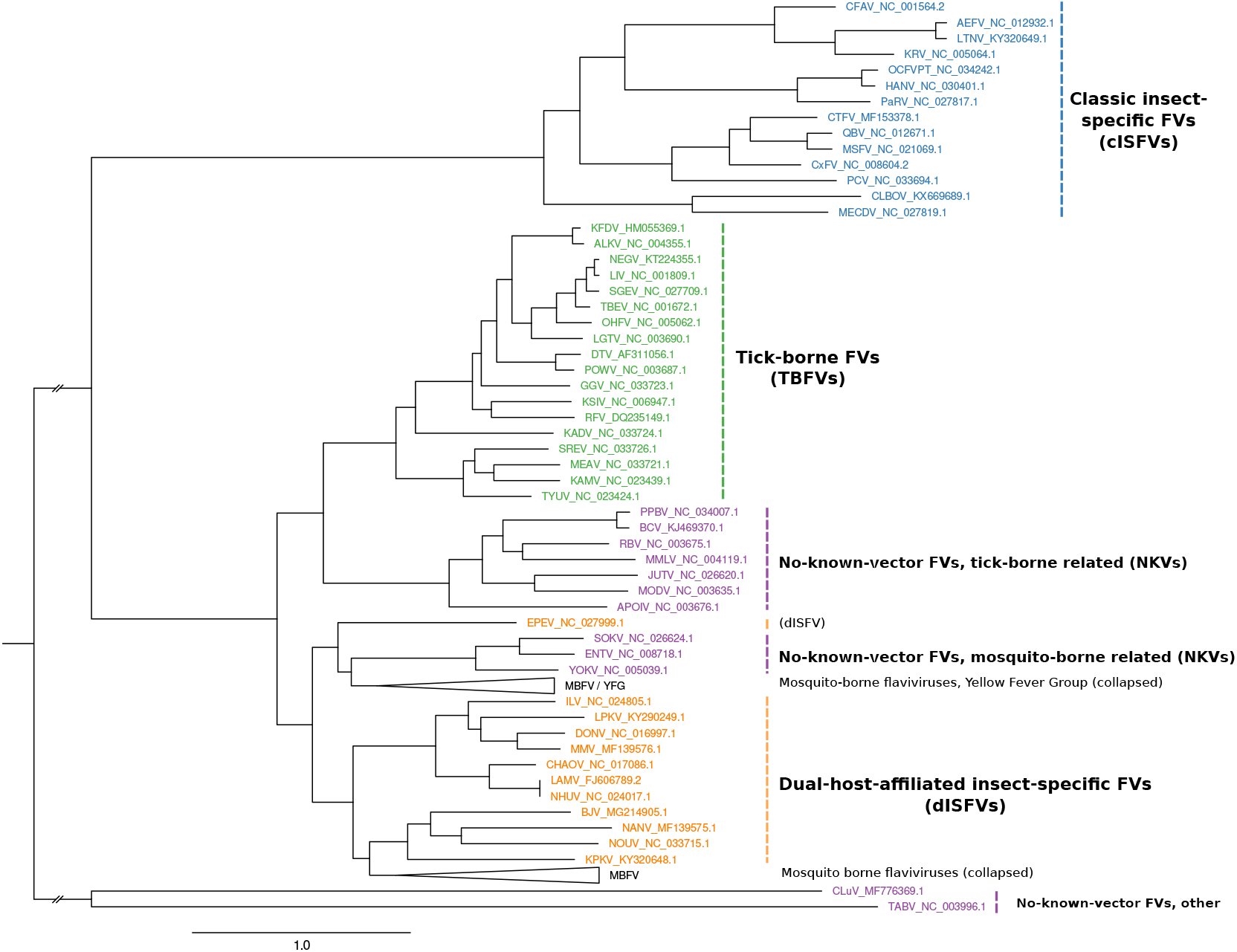
Maximum-likelihood phylogenetic tree of the genus *Flavivirus*, highlighting the major groups ISFVs (blue), dISFVs(orange), TBFVs (green), and NKVs (magenta). The MBFV Yellow Fever virus group (YFG) and the main MBFV branch were not covered in this study and are both collapsed. The tree has been computed from a `MAFFT` alignment of complete polyprotein amino acid sequences with `iq-tree`. Figure rendered with `FigTree`.

TBFVs form a monophyletic group consisting of a single serocomplex, although pathology and clinical manifestations vary among different viruses. They comprise more than a dozen of recognized species and separate into three groups: Mammalian tick-borne flaviviruses (M-TBFV), seabird tick-borne flaviviruses (S-TBFV) and the Kadam virus group. See [7] for a comprehensive review.

ISFVs naturally infect hematophagous *Diptera* and are typically divided into two groups [8]: Classical insect-specific flaviviruses (cISFVs) naturally infect mosquitoes and excursively replicate in mosquito cells *in vitro*. They form a phylogenetically distinct clade among known flaviviruses, appearing at the root of the MBFV, TBFV and NKV branches. The cISFV group separates into two clades, one associated with *Aedes* spp. mosquitoes and the other associated with *Culex* spp. mosquitoes, respectively [9]. They lack the ability to infect vertebrates and to replicate in vertebrate cell lines and have not been in the research spotlight until very recently. The second group is comprised of arbovirus-related or dual-host affiliated insect-specific flaviviruses (dISFVs), which represent a non-monophyletic group which is phylogenetically and antigenically related to mosquito/vertebrate flaviviruses, although they do not appear to infect vertebrate cells [10]. Insect-specific viruses play a crucial role in the mosquito microbiome and have been shown to modulate the replication of other arboviruses [11]. In this line, they are currently considered as biological control agents and vaccine platforms [12].

NKVs represent an ecologically and phylogenetically diverse set of viruses which have been isolated exclusively from vertebrates (mainly bats and rodents), without evidence for transmission by arthropod vectors. They form a non-monophyletic group among flaviviruses and are typically divided into bat- (B-NKV) and rodent-associated (R-NKV) groups, see Table A1. B-NKVs can be further separated into Entebbe virus group, which is phylogenetically closer to MBFVs, and Rio Bravo virus group, which is a sister clade to TBFVs. Species in the R-NKV group form the Modoc virus group, which is phylogenetically close to the B-NKV Rio Bravo group [13]. While NKVs are poorly characterized they represent a valuable resource to study evolutionary traits related to host-switch capacity mediated by conserved genomic elements.

### 1.1. Conserved RNA structures mediate pathogenesis

Conserved RNA structures in the untranslated regions (UTRs) of RNA viruses are of particular interest because they mediate the viral life cycle by promoting or enhancing replication, as proposed for elements in both 5’UTRs [14] and 3’UTRs [15–18]. Mosquito/vertebrate viruses must operate efficiently in vectors and hosts, phylogenetically distinct organisms with different cellular machineries. This requires a high level of flexibility of viral regulatory elements to evade various antiviral response strategies while assuring proper replication conditions required for maintaining a stable quasispecies population. To achieve this resilience in host adaptation, RNA duplication strategies have been proposed as an evolutionary trait for MBFVs [19]. Tandem RNA structures within DENV 3’UTR are under different selective pressures in alternating hosts, suggesting the idea that duplicated RNA structures differentially evolved to accommodate specific functions in the two hosts [20]. Likewise, there is evidence for evolutionary pressure on maintaining the primary sequence of parts of duplicated RNA elements, as recently shown for flaviviral dumbbell (DB) elements in the context of finding a biophysical model for explaining a possible route for ZIKV-induced neurotropism [21].

Viral RNA genomes are different from procaryotic and eucaryotic mRNA. In addition to coding for and regulating the viral machinery, viral genomic RNA (gRNA) exhibits functional regions that act upon different stages of the viral life cycle. The 10-12kB flaviviral gRNA is capped, but non-polyadenylated and encodes a single open reading frame (ORF). Upon translation, a polyprotein is produced, which is then cleaved by viral and cellular enzymes into structural and non-structural proteins [22]. The ORF is flanked by highly structured untranslated regions (UTRs), which contain evolutionary conserved RNA elements that are crucially related to regulation of the viral life cycle, thereby inducing processes such as genome circularization, viral replication and packaging [23–25].

Upon flavivirus infection, accumulation of both gRNA as well as viral long non-coding RNAs (lncRNAs) is observed. These lncRNAs, which have been referred to as subgenomic flaviviral RNAs (sfRNAs) [26] are stable decay intermediates produced by exploiting the host’s mRNA degradation machinery [27] and are associated with viral replication, pathogenesis and cytopathicity [28,29]. The production of sfRNA is induced by partial degradation of viral gRNA by the 5’-3’ exoribonuclease Xrn1, an enzyme associated with the cell’s RNA turnover machinery [30,31]. Mechanistically, sfRNAs are generated by stalling Xrn1 at conserved structural elements in the viral 3’UTR, termed xrRNA (exoribonuclease-resistant RNA elements). These structures efficiently stall Xrn1 from progressing through from the 5’ direction, thus protecting the downstream RNA from degradation, while pass-through in the 3’-5’ direction, as required for viral RNA-dependent RNA-polymerase is still possible [32]. In particular, different types of stem-loop (SL) and dumbbell (DB) elements found in many MBFVs and TBFVs have been related to quantitative protection of downstream virus RNA against exoribonuclease degradation [33].

Xrn1 stalling results in dysregulation of cellular function with the aim of promoting viral infections. In this regard, functions of sfRNA in modulating cellular mRNA decay and RNAi pathways [34] as well as modulating anti-viral interferon response [35,36] have been reported.

The genomic architecture of flaviviruses has been extensively studied to understand the molecular principles required for sfRNA production. Chemical and enzymatic probing methods [37], together with x-ray crystallography revealed the 3’UTR structure of the MBFVs WNV [38], YFV [39], DENV [40], Murray Valley encephalitis virus (MVEV) [41], ZIKV [42] and recently different species of the TBFV and NKV groups [33], highlighting the possibility that exoribonuclease resistance might be a pervasive mechanism of the viral world. Interestingly, several conserved RNA structural elements in viral 3’UTRs have been predicted in our group [43–47], some of which have later been attributed to xrRNA functionality [26]. To further expand the set of potential xrRNAs, we report here on a comparative genomics survey aimed at characterization of evolutionary conserved RNA structures in flavivirus 3’UTRs, focusing on TBFVs and the hitherto understudied groups of ISFVs and NKVs. A detailed study on the evolutionary traits of conserved RNAs in MBFV 3’UTRs will be published elsewhere.

## 2. Materials and Methods

Viral genome data for the present study were obtained from the public National Center for Biotechnology Information `(NCBI) refseq` (https://www.ncbi.nlm.nih.gov/refseq/) and `genbank` (https://www.ncbi.nlm.nih.gov/genbank/) databases on 28 May 2018. We downloaded all complete viral genomes under taxonomy ID 11051 (genus *Flavivirus*) and filtered for TBFV, ISFV and NKV species listed in table A1. Whenever `refseq` annotation was not available for a species, we selected the longest complete genome from the `genbank` set as representative sequence. In total, the data set is comprised of 86 ISFV, 275 TBFV and 27 NKV isolates, respectively. The number of isolates with available 3’UTR sequence data per species varies between 1 and 167.

### 2.1. Phylogeny reconstruction

The polyprotein/coding sequence (CDS) regions of most flaviviruses can be aligned consistently, however, UTRs typically show large variance both in length and sequence composition, rendering them ill-suited for phylogeny reconstruction. A phylogeny of all members of the genus *Flavivirus* (Fig. 1) was therefore reconstructed via a multiple sequence alignment (MSA) of the nucleotide sequences of the CDS regions only. The MSA was computed with `MAFFT`[48] and subsequent maximum-likelihood tree reconstruction was performed using `iq-tree`[49] using the GTR+F+R7 substitution model.

### 2.2. Structural homology search with covariance models

The present study is centered around structural homology of RNA elements among phylogenetically narrow subgroups. A straightforward approach to finding novel homologous RNA structures is to search RNA sequence databases with Covariance Models (CMs), i.e. statistical models of RNA structure that extend classic Hidden-Markov-Models (HMMs) to simultaneously represent sequence and secondary structure. CMs, as implemented in the infernal package[50] allow for rapid screening of large RNA sequence databases to find even weakly conserved sequence-only or structurally homologous RNAs. We have recently applied this approach to identify novel telomerase RNAs in *Saccharomycetes* [51].

Here, structural multiple sequence alignments of the viral 3’UTR sequences were generated with `locARNA` [52] and CMs were built for known or experimentally verified xrRNAs [33]. All 3’UTR sequences were then screened and novel candidate sequences were added to perform iterative refinement until convergence. Weak sequence conservation of putative xrRNA elements resulted in initially fragile results, indicating that `infernal default` parameters are typically not optimal. Adjusting parameters, in particular disabling both heuristic filtering and local end detection, however, allowed our CMs to find homologs with strongly conserved secondary structures in presence of large sequence deviation from the original sequence the CM was built from. Likewise, `cmsearch` E-values turned out unsuitable for assessing hit quality in case of major sequence divergence. We therefore employed a cutoff approach, requiring a hit to form at least 75% of all base pairs listed in the CM in order to be considered significant.

### 2.3. De novo discovery of conserved RNA elements

Beside characterization of RNAs with homology to known structurally conserved elements, we aimed at identifying novel elements, considering both thermodynamic stability and sequence covariation as evolutionary traits. In this line, `locARNA`-generated structural alignments of full UTR sequences were cut manually into blocks corresponding to conserved secondary structures. Alternatively, we employed `RNALalifold` from the `ViennaRNA` package [53] to compute locally stable secondary structures for aligned UTR sequences. A CM was built for each structure and searched against all flavivirus 3’UTRs, keeping only CMs that scored well multiple times per UTR. The rationale here is that the occurrence of multiple copies hints towards a possible functional role of a structural element, given that the ability of two or more independently evolving sequences to form a common structure is unlikely.

The above approach is implemented as a set of custom Perl and Python scripts for semi-automatic characterization and annotation of conserved RNAs in viral UTR sequences. Internally, these scripts build on the `ViennaRNA` scripting language interface for thermodynamics calculations, the `ViennaNGS` [54] suite for extraction of genomic loci, the `RNAaliSplit` package [55] for splitting alignments into subparts with common consensus structures (i.e. common structures formed by all individual sequences), `R2R` [56] for visualization, and the `ETE3` framework [57] for tree annotation and visualization.

## 3. Results

Several flaviviruses have previously been studied in great detail, yielding a varied landscape of repeated RNA sequence and structure elements within the 3’UTRs of these viruses, which are likely to have evolved from numerous duplications [19,58,59]. Many of these studies relied on single sequence predictions, which resulted in a good understanding of both structure and genomic position of conserved elements in individual species. A unified picture of homologous RNAs within the 3’UTRs of flaviviruses, however, has not been available.

The comparative approach applied in the present study outperforms single sequence predictions by considering consensus structures formed by all sequences. This allows us not only to confirm previously described RNA structures but also to elucidate hitherto unrecognized tandem repeats in many species. In this line our results can help in understanding the complex evolution of flavivirus 3’UTRs.

### 3.1. Construction of seed alignments

Based on recent experimental evidence for the existence of xrRNAs in TBFVs, ISFVs and NKVs, and previously characterized conserved RNA elements in flaviviral 3’UTRs, we built seed alignments for initial CMs, which were then refined iteratively, i.e., subjected to multiple rounds of screening and incorporation of best hits into the CM. Likewise, candidate structures from `RNALalifold` calculations were used as seeds for identification of conserved RNA structures. Fig. 2 shows an overview of refined consensus structures for each ecologic group of flaviviruses analyzed here.

**Figure 2.**
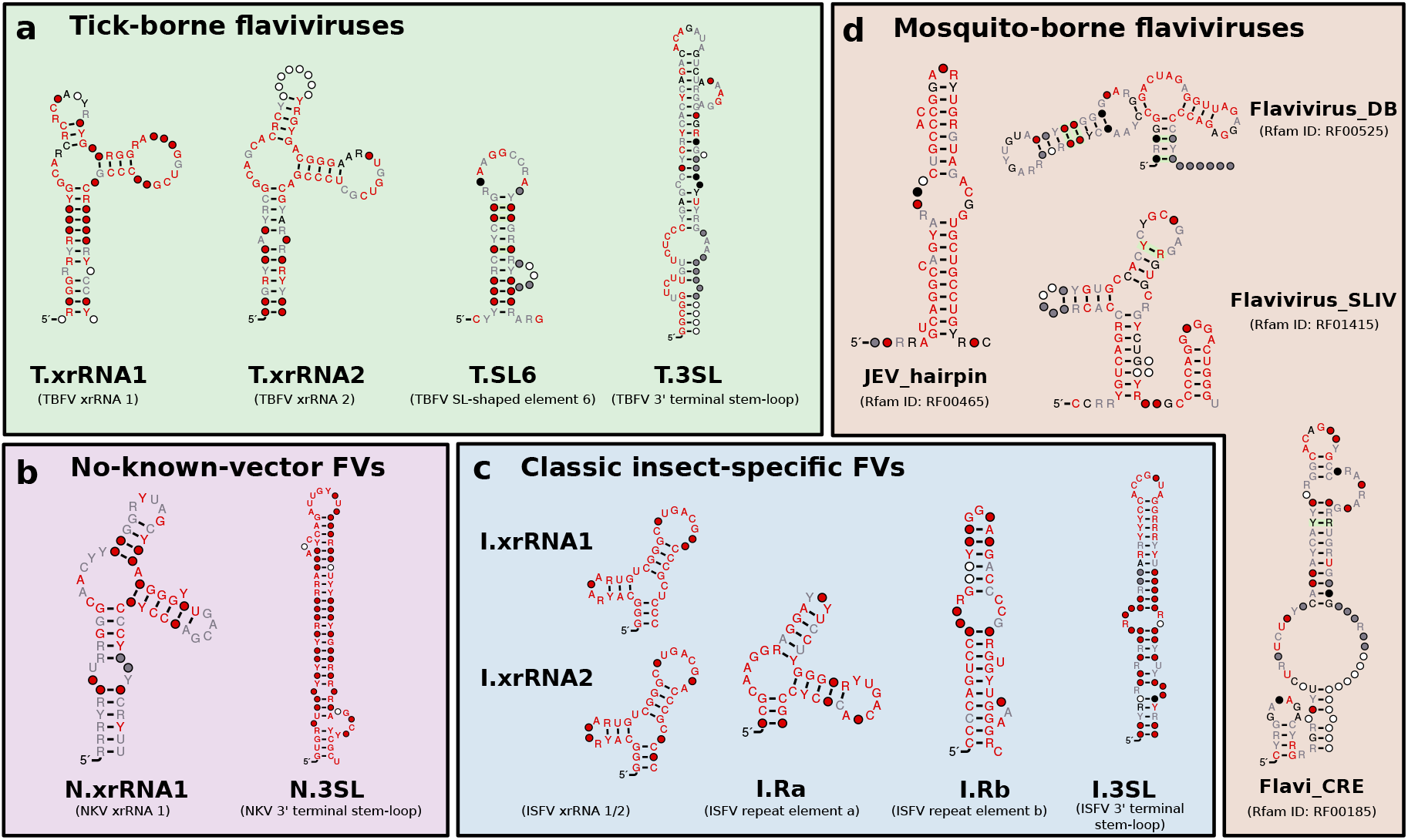
Overview of consensus structures of all CMs used for the annotation of flavivirus 3’UTRs. TBFV, ISFV, and NKV elements were refined from published experimental data (T.xrRNA1/2, I.xrRNA1/2, N.xrRNA) or identified computationally (T.SL6, I.Ra, I.Rb as well as all 3’-terminal stem-loop structures). MBFV elements were obtained from Rfam. Throughout this paper, all CMs are referred to by the name written in bold. References to xrRNA-like structures refer to the generalized xrRNA CM (Section 3.6).

The four ecologic groups of flaviviruses show a varied 3’UTR architecture, however, the terminal 3’ stem-loop structure (3’SL, also referred to as 3’ long stable hairpin, 3’LSH) has been shown to be associated with panhandle-formation during virus replication and is therefore present in the terminal region of all flaviviruses [14]. The element is listed in Rfam as RF00185 (Flavivirus 3’UTR cis-acting replication element, Flavi_CRE) and we could use it to consistently identify terminal regions within 3’UTRs. Absence of this element from a UTR sequence is indicative of incomplete or truncated sequence data. The underlying sequences generally form a stable stem-loop structure upon structural alignment and single sequence folding. We built individual 3’SL seed alignments and CMs for each ecologic group, termed T.3SL, N.3SL and I.3SL, respectively.

### 3.2. Tick-borne flaviviruses

MacFadden *et al*. [33] suggested two different exoribonuclease-resistant structures in TBFV 3’UTRs. We used the proposed sequences from TBEV, POWV, Karshi virus (KSIV), Langat virus (LGTV), Louping ill virus (LIV), Omsk hemorrhagic fever virus (OHFV) and Alkhumra hemorrhagic fever virus (ALKV) as templates for a set of initial structural alignments and CMs. These models were then employed to search for high confidence hits within all TBFV 3’UTRs to construct seed alignments of the two exoribonuclease resistant structures in TBFVs, termed T.xrRNA1 and T.xrRNA2. These models allowed us to construct highly specific CMs for both TBFV xrRNAs, which were subsequently used to annotate xrRNA instances in already studied and previously unstudied TBFV species (Fig. 3 a).

**Figure 3.**
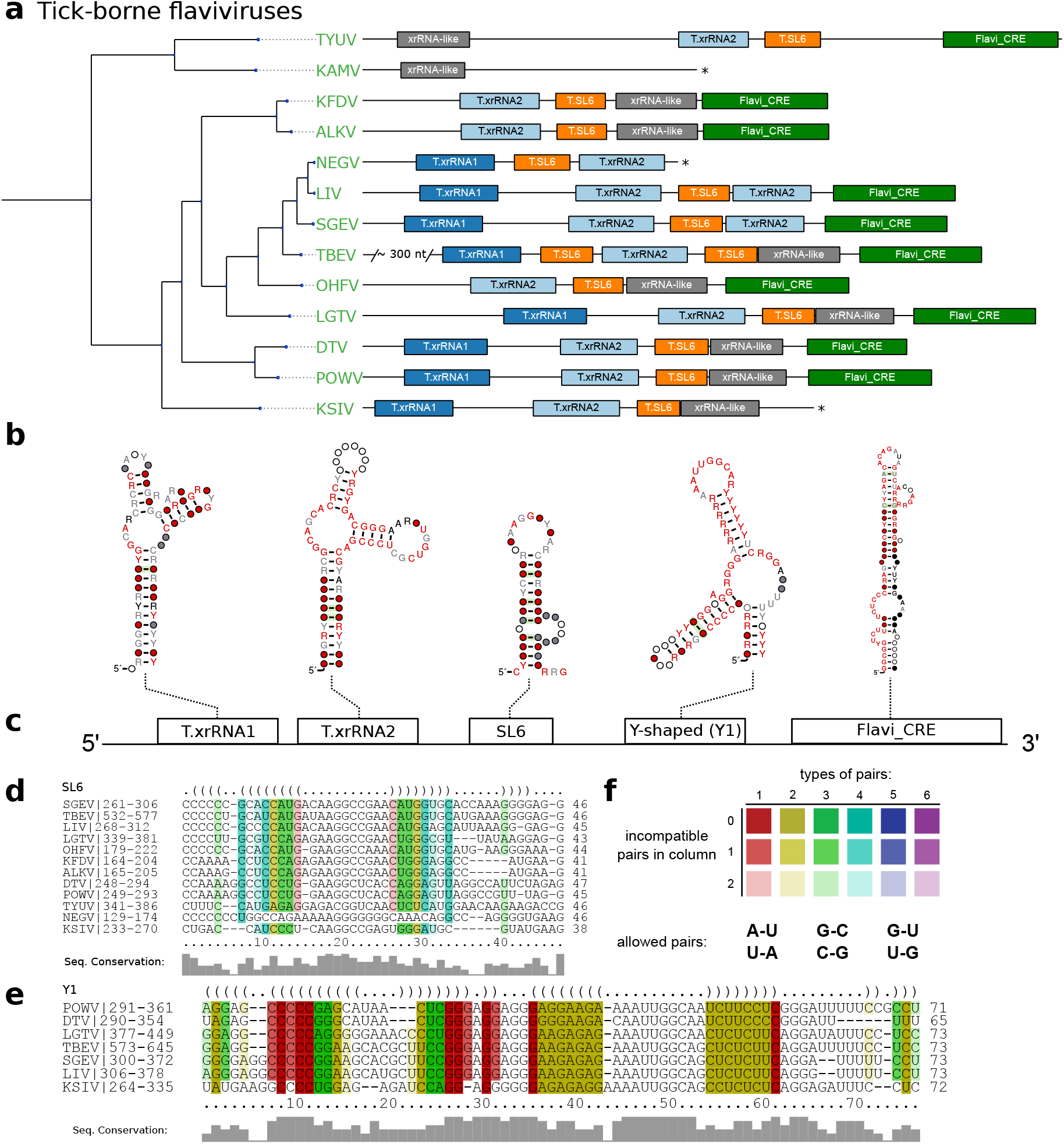
a Annotated 3’UTRs of TBFVs. The phylogenetic tree on the left has been computed from complete coding sequence nucleotide alignments and corresponds to the TBFV subtree in Fig. 1. For each species with available 3’UTR sequence a sketch of the 3’UTR is drawn to scale next to the leaves of the tree. Colored boxes represent conserved RNA structural elements. Identifiers within the boxes indicate the CM which was used to infer homology at this position. Asterisks indicate incomplete 3’UTR sequences. Species without available 3’UTR are not shown. b Consensus structure plots of CM hits as calculated by *mlocarna*. c Schematic depiction of the common structural architecture of TBFV 3’UTRs. d,e Structural alignments of elements SL6 and Y1. f RNAalifold coloring scheme for paired columns in alignments. Colors indicate the number of basepair combinations found in pair of columns. Fainter colors indicate that some sequences cannot form a base pair.

The full structural alignment of the 3’UTRs of selected tick-borne species moreover suggests a short stem-loop element in several species, which is characterized by high sequence heterogeneity but heavily conserved structure supported by multiple covariations. Evidence for this element, termed stem-loop 6 (SL6), has been reported earlier for at least TBEV, LGTV and OHFV [58,59]. We kept this nomenclature and identified the exact position in each TBFV 3’UTR (Fig. 3d).

Our data further shows that both TBFV xrRNA CMs (Fig. 2 a), as well as NKV xrRNA CMs (Fig. 2 b and Sect. 3.5) consistently yield plausible hits with a high degree of structure conservation immediately upstream of the strongly conserved terminal stem-loop element. Existence of a Y-shaped element (termed Y1) and putative similarity to NKVs has been proposed earlier based on single sequence structure predictions [58]. Structural `locARNA` alignment and subsequent `RNAalifold` consensus structure prediction indicates strong secondary structure conservation with frequent structure-conserving sequence covariations. Taken together, this suggests good evidence that respective regions in TBFVs harbor a putatively structured and functional xrRNA-like RNA (Y1, Fig. 3 d).

Despite the differences in length and sequence composition, the 3’UTRs of most species in the TBFV group share a common architecture. Similar to MBFV SL-elements [19], two copies of xrRNAs are found in almost every species of this ecologic group, generally succeeded by one instance of SL6 and Y1. Likewise, the terminal 3’ stem loop is conserved in all TBFVs and can be reliably annotated by both our CM, T.3SL, and the Rfam Flavi_CRE model, which is used in Fig. 3. Among all investigated species, only ALKV, OHFV and Kyasanur forest disease virus (KFDV) do not have a copy of xrRNA1, indicating that these viruses may have previously lost this element. Conversely, the two seabird-associated TBFVs with available 3’UTR data, Tyuleniy virus (TYUV) and Kama virus (KAMV) do not fit into this general scheme. Likewise, we were not able to annotate additional homologous or conserved structures with any CM used in this screen in the variable region of the 3’UTR of TBEV [5], despite the substantially longer UTR (+300 nts).

### 3.3. Classic insect-specific flaviviruses

Classic insect-specific flaviviruses present diverged 3’UTR architectures, which likely result from the association of different species to *Aedes* spp. and *Culex* spp. vectors, respectively, which are also reflected by clade separation in the ISFV phylogenetic tree. Previous studies employed single sequence predictions to propose a varied set of homologous RNA structures in combination with an unusually large number of duplicated sequence signals [59]. Recent experimental evidence, however, suggests the presence of xrRNAs that have a similar fold to those known from MBFVs in cISFVs. Consequently, we set out to independently characterize conserved RNA elements for different subclades.

#### 3.3.1. Exoribonuclease-resistant RNAs in Aedes-associated cISFVs

MacFadden *et al*. [33] utilized SHAPE structure probing to report the presence of two exoribonuclease-resistant RNAs in Cell fusing agent virus (CFAV), and provided evidence for a duplicated set of homologous structures in Aedes flavivirus (AEFV) and Kamiti river virus (KRV). We constructed initial alignments from the reported sequences in this clade in *Aedes* spp. associated viruses, resulting in two seed alignments, termed I.xrRNA1 and I.xrRNA2 (Fig. 2 c). For both elements, seed CMs were iteratively built from structural `locARNA` alignments. Minor manual adjustments to the alignments were required here, since the predicted consensus structures diverged slightly from the published SHAPE-guided prediction. Both models were then employed to search for additional high confidence hits within other isolates of CFAV, AEFV and KRV which were subsequently added to the seed alignments.

Screening the entire set of flavivirus 3’UTRs revealed that both ISFV xrRNA elements, I.xrRNA1 and I.xrRNA2, are only found in CFAV, AEFV and KRV, i.e., species initially used for the construction of the respective CMs. Furthermore, also in terms of pure structural conservation, no reliable hits in any other ISFV species could be obtained with any of these CMs (Fig. 4 a). This suggests that both ISFV xrRNAs may represent a specialized class of xrRNA elements only present in CFAV, AEFV and KRV. The 3’UTR of KRV is unique among all known flaviviruses because it harbors an additional copy of the terminal 3’ stem-loop element 600 nts upstream of the actual 3’-terminus, supporting previous reports that the KRV 3’UTR has undergone a full duplication during its evolution [60].

**Figure 4.**
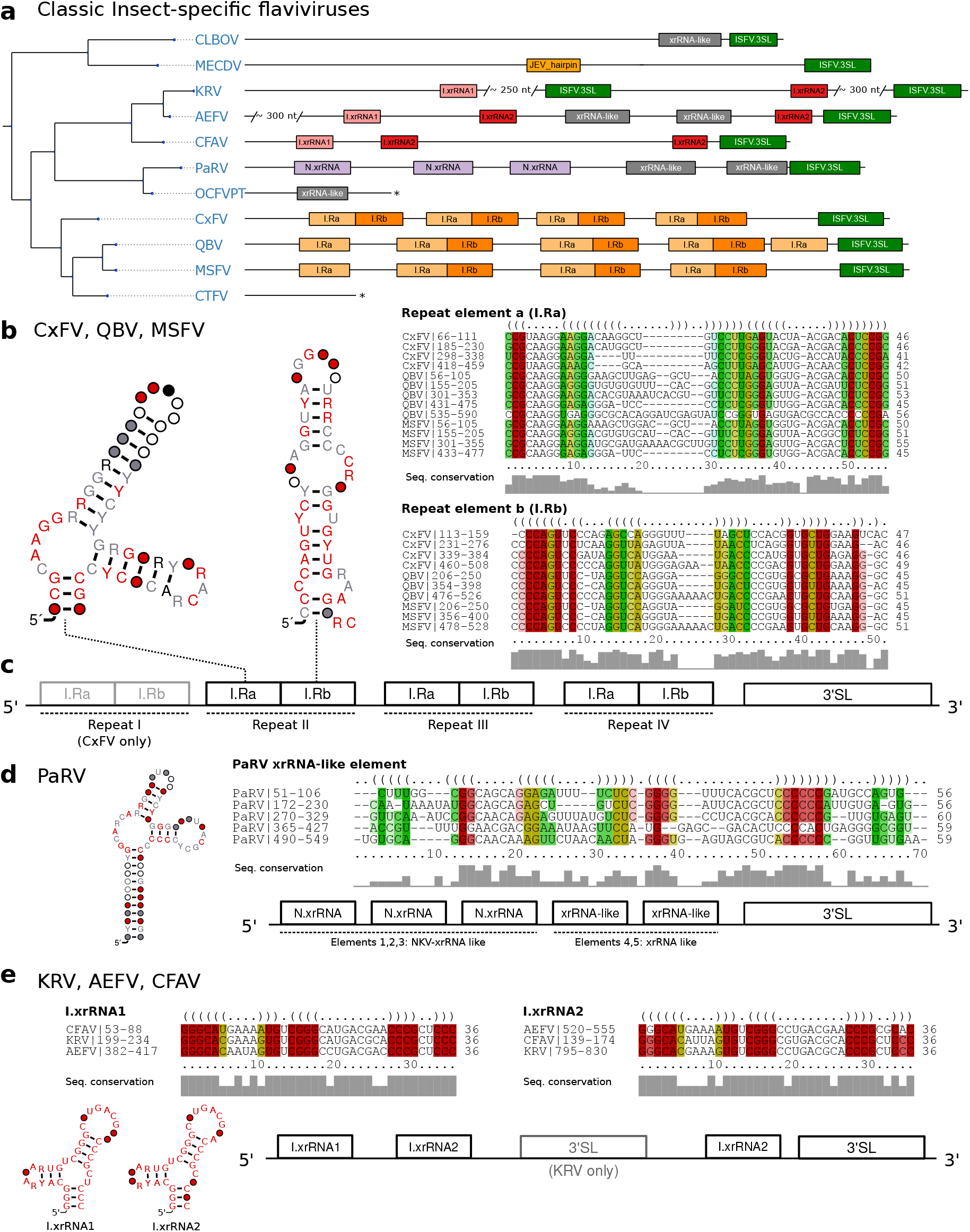
a Annotated Tree of cISFV 3’UTRs. Asterisks denote incomplete 3’UTR sequences. Species without available 3’UTR are not shown. b Consensus secondary structure plots and structural alignments of CM hits of Repeat a/b elements in CxFV, QBV, and MSFV. c Schematic of the common architecture of CxFV, QBV, and MSFV. Element I-IV refers to the respective repeat of elements. d Consensus structure plot and structural alignment of all CM hits of xrRNA-like elements in PaRV. e Proposed 3’UTR architecture of KRV, AEFV, and CFAV with consensus structure plots and structural alignments of I.xrRNA1 and I.xrRNA2.

#### 3.3.2. Conserved structures in *Culex*-associated cISFVs

The second distinct clade of cISFVs includes Culex flavivirus (CxFV), Quang Binh virus (QBV), Mosquito flavivirus (MSFV), Palm Creek virus (PCV), Culex theileri flavivirus (CTFV) as well as a few other species with only partial genome sequence availability [9] and is associated with *Culex* spp. vectors. An interesting observation in this clade is that no other CM from any of the four ecologic flavivirus groups shows a hit, not even with remote sequence or structure conservation. We therefore set out to produce a a high quality structural alignment of the complete 3’UTRs of CxFV, QBV and MSFV. Consensus structure folding of the full alignment revealed each species to harbor 3-4 repeats of two highly conserved elements supported by multiple co-varying base pairs (Fig. 4 b, c). We termed these “Repeat element a/b”, respectively (Ra and Rb). Both elements, while strongly conserving their folds, show highly variable loop regions as well as weak sequence conservation in the case of the Ra element. Structure conservation and occurrence in multiple copies, as typically seen with other exoribonuclease-stalling elements, hints towards possible functional importance. These results complement earlier reports of sequence repeats in the 3’UTR of CxFV [61] with the identification of a quadruplicated pair of conserved structures.

#### 3.3.3. Diverged 3’UTR architecture in many cISFVs

Interestingly, a screen of all available CMs in Parramatta River virus (PaRV) revealed five xrRNA-like elements (Fig. 4 a), with elements 1-3 sharing sequence and structure properties with NKV xrRNAs (Section 3.5), while elements 4 and 5 only conserve N.xrRNAs structure. All five hits can be structurally aligned into a consistent consensus structure (Fig. 4 d), despite the overall weak sequence consensus.

Conversely, the 3’UTRs of Calbertado virus (CLBOV) and Mercadeo virus (MECDV) appear structurally different from the other cISFVs. A general lack of characteristic CM hits lets these species appear more like an outgroup among cISFVs. In particular, we could only find significant hits for the omnipresent terminal 3’stem-loop structure, a putative xrRNA-like element in CLBOV and a single instance of a structure homologous to the Rfam model RF00465 (Japanese encephalitis virus hairpin structure) in MECDV. Still, limited availability of 3’UTR sequence data renders the characterization of conserved elements and interpretation difficult here.

Our data suggests that the 3’UTRs of cISFVs, in contrast to TBFVs (Section 3.2 and dISFVs (Section 3.4), do not appear to have a consistent architectural organization. In agreement with the cISFV phylogenetic subtree (Fig. 1) we constitute three diverged groups with common 3’UTR organization that conform to their respective sub-clades: (i) CFAV-AEFV-KRV, each with two instances of xrRNAs, (ii) CxFV-QBV-MSFV with 3-4 copies of I.Ra/I.Rb elements and (iii) PaRV with 4-5 copies of xrRNA like structures. Although no full 3’UTR sequences are available for the phylogenetically closest relatives of PaRV, HANV and OCFVPT, an xrRNA-like element in the small available fragment (syntenic to PaRV UTR) of OCFVPT 3’UTR suggests that both viruses might be organized in a similar manner, as supported by earlier reports that these viruses should be classified within the same species [1]. For CLBOV and MECDV, no clear pattern of conserved elements can be identified with our CMs. Both viruses either employ an entirely different class of elements or might not require capability for exoribonuclease stalling at all. The only element shared universally among all cISFVs is the 3’-terminal stem-loop, although cISFVs seem to diverge from other flaviviruses here, indicated by the inability of Rfam model RF00185 (Flavivirus CRE) to reliably annotate any cISFV 3’UTR.

### 3.4. Dual-host affiliated insect-specific flaviviruses

Isolated almost exclusively from mosquitoes, dISFVs do not seem to infect vertebrate cells, despite their phylogenetic proximity to MBFVs (Fig. 1). This association is reflected by good hits of the Rfam covariance models RF00525 (Flavivirus DB element) and RF00465 (Japanese encephalitis virus hairpin structure) in all dISFV isolates studied here (Fig. 5). Interestingly, we could not find evidence for any sequences or structures homologous to tick-borne or other insect-specific flaviviruses. In this line, our data is in good agreement with the phylogenetic location of these viruses, which share ancestral roots with MBFVs [10].

**Figure 5.**
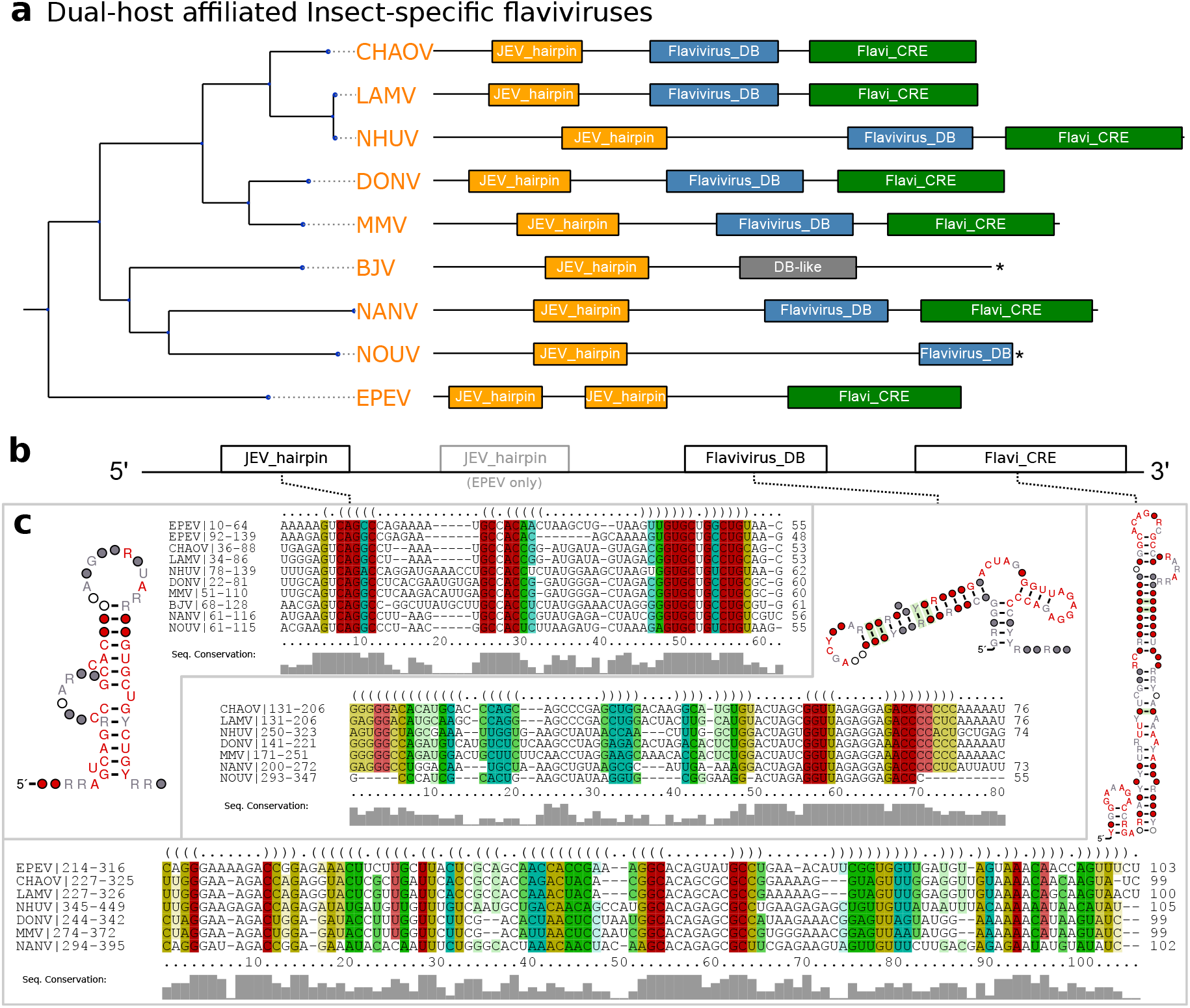
a Annotated Tree of dISFV 3’UTRs. Asterisks denote incomplete 3’UTR sequences. Species without available 3’UTR are not shown. b Schematic Architecture of the dISFV 3’UTR. c Structural alignments and consensus structure plots of dISFV elements.

An unusual species within this group is Ecuador Paraiso Escondido virus (EPEV), which has been isolated from New World sandflies and has been classified as insect-specific virus. EPEV phylogenetically appears at the root of the Entebbe bat virus group (ENTVG), a clade comprised of the three NKVs Entebbe bat virus (ENTV), Sokoluk virus (SOKV) and Yokose virus (YOKV). While all of these viruses contain homologs of conserved stem-loop (SL) and dumbbell (DB) elements found in MBFVs, ENTVG species may have lost their vector dependence [1].

### 3.5. No-known-vector flaviviruses

Rather than forming a monophyletic group, the no-known-vector flaviviruses can be separated into two distinct lineages, which are closely related to either TBFVs or MBFVs, respectively (Fig. 1). Two additional NKVs, Tamana bat virus (TABV) and Cyclopterus lumpus virus (CLuV), are phylogenetically distant and serve as an outgroup to all flaviviruses. In analogy to the procedure outlined above for TBFVs (Section 3.2) and ISFVs (Section 3.3), we built a CM for experimentally verified xrRNAs in tick-borne related NKVs, termed N.xrRNA.

We found multiple hits of this CM at various loci within the 3’UTRs of tick-borne related NKVs, indicating that these species, in contrast to TBFVs, do not conserve a common 3’UTR architecture (Fig. 6 a). Surprisingly, we could identify several high-quality hits of the Rfam model RF00525 (Flavivirus DB element), an element typically found in MBFVs, in Rio Bravo virus (RBV), Montana myotis leukoencephalitis virus (MMLV) and Modoc virus (MODV). This is in so far remarkable as there is no evidence for conservation of this element in TBFVs, which phylogenetically cluster with this clade of NKVs. This element might have been introduced by an ancestral recombination event. Alternatively, conservation of an MBFV element in NKVs might be indicative of an association to an unknown vector, in agreement with the hypothesis that vector specificity is mediated by characteristic 3’UTR elements [19].

**Figure 6.**
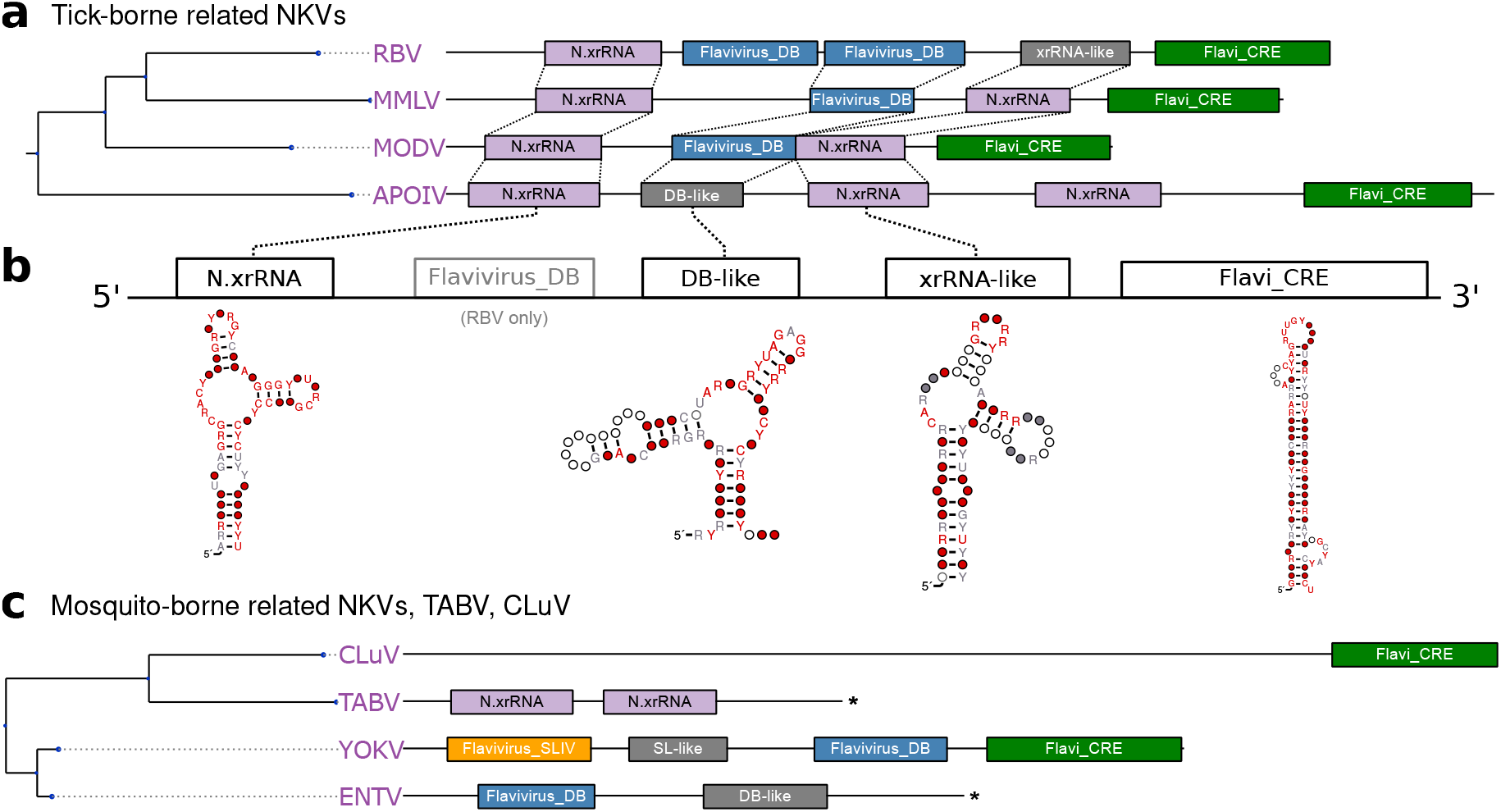
a,c Annotated 3’UTRs of NKVs. Asterisks denote incomplete 3’UTR sequences. b Schematic of TBFV-associated NKV-FV UTR architecture with consensus structures of NKV structure elements.

Conversely, there seems to be no generally conserved 3’UTR architecture among members of the mosquito-borne related NKVs (Fig. 6 c). While sequence data has not been available for Sokoluk virus (SOKV), we could annotate typical MBFV elements in the next relatives Entebbe bat virus (ENTV) and Yokose virus (YOKV), as proposed previously [32].

### 3.6. A generalized xrRNA structure

Earlier work suggested that xrRNAs from TBFVs and tick-borne related NKVs fall into a more general structural class of xrRNAs [33]. Following this line of reasoning, we investigated whether all high confidence hits obtained with our TBFV and NKV CMs could be assembled into one coherent CM that conserves the xrRNA-typical fold. A further advantage of a generalized CM would be higher sensitivity, allowing for identification of common features and eventually lead to annotation of previously unannotated xrRNAs.

Structural alignment and consensus structure prediction revealed all high confidence hits to fold into a common secondary structure (Fig. 7 a, b). While most of the consensus structure is characterized by low sequence conservation, stem 3 (S3) and loop 1 (L1) show medium to high degree of sequence conservation. The length of all stems is well conserved, although both major loop regions L2 and L3 show large fluctuations, with the length of L3 being de facto constant and L2 showing a high degree of flexibility.

**Figure 7.**
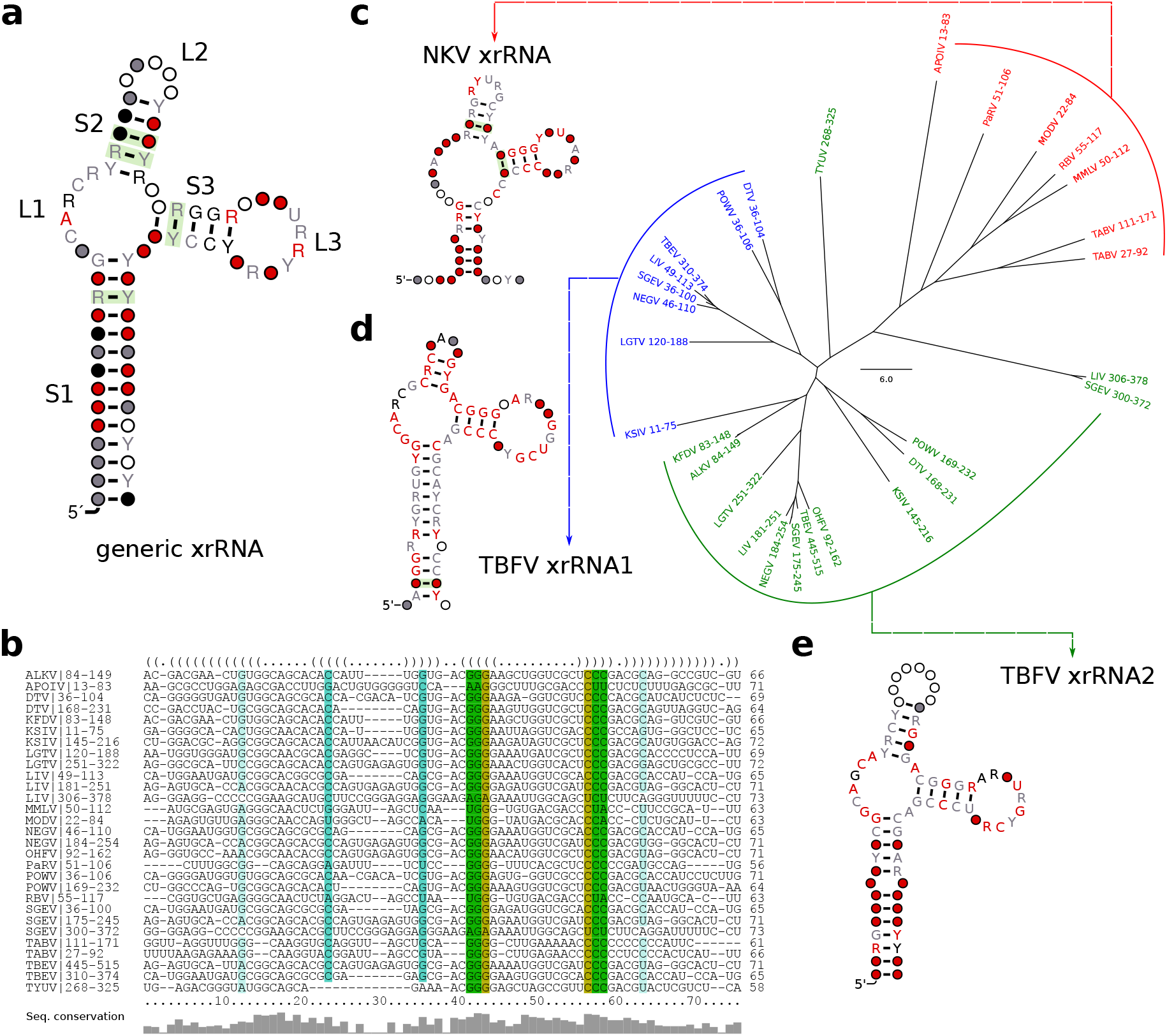
Generalized structure of all high confidence (cmsearch evalue < 10^−5^) hits of T.xrRNA1, T.xrRNA2 and N.xrRNA. **a** Consensus structure prediction and **b** structural alignment of all high confidence hits. **c-e** Neighbor-joining tree of all high confidence hits of N.xrRNA (c), T.xrRNA1 (**d**), and T.xrRNA2 (**e**). Leaves are grouped and colored by the CM used for annotation, coordinates correspond to the position in the respective 3’UTR. For each group a separate structural alignment was computed, the consensus structures are shown.

We further investigated whether any high confidence hits from I.xrRNA1/2 in cISFVs could be aligned to the generalized xrRNA model. Although both cISFV xrRNAs (Fig. 2 c) bear some similarity to the generalized model, in particular to S3 and L3, we were not able to build a common alignment or consensus structure. Despite seemingly similar shape, our data also suggests that cISFV xrRNAs form a separate xrRNA subclass, unrelated to MBFV xrRNAs. In particular, we could not obtain hits of Rfam CMs (which can be seen as representatives of MBFV elements) in cISFVs, nor could we confirm any hits of cISFV-specific elements in MBFVs.

In addition to learning xrRNA features, a more generalized CM enabled us to detect xrRNA-like structures (indicated as such in all annotation plots), that could not be found previously.

## 4. Discussion

Mediated gRNA decay in the form of exoribonuclease resistance seems to be a pervasive strategy employed by viruses to circumvent host immune responses. Evidence of sfRNA production following incomplete Xrn1 degradation has not only been observed in different members of the *Flavivirus* genus [62], but also in other species of the *Flaviviridae* family, however, with major differences in xrRNA structure and sfRNA characteristics. While MBFV produce a 300-500nt sfRNA that corresponds to degradation products of the gRNA 3’UTR, hepaciviruses and pestiviruses produce a long subgenomic RNA whose 5’ end is located within the first 130nt of the viral gRNA [63].

Moreover, recent studies have identified xrRNA functionality in several phylogenetically distant RNA viruses, such as animal-infecting, segmented viruses of the *Bunyaviridae* and *Arenavividae* [64] families, as well as plant-infecting viruses of the *Tombusuviridae* and *Luteoviridae* families [65,66]. The interesting question whether exoribonucleases other than Xrn1 would be blocked as well has recently been answered. MacFadden *et al*. [33] could show that both RNAse J1 and Dxo1 are stalled by MBFV xrRNAs, thereby demonstrating the general nature of this structure-induced blocking mechanism. These novel findings, together with previous knowledge of Xrn1 stalling in segmented plant viruses [67,68] provide evidence for a convergent evolution scenario where xrRNAs depend on a specific folded RNA structure and form a distinct class of functional RNAs.

Repeated RNA elements appear to be a hallmark of flavivirus 3’UTR architecture. While there seems to be a plethora of conserved structure classes, our data emphasizes the consistent trend that these elements typically do not occur as single copies. Rather, duplicate or even multiple occurrences of these elements hint towards functional relevance. This is further underlined by the evolutionary conservation of both patterns and elements among different species. Since, exoribonuclease stalling is presumably never perfect, it makes sense that viruses might employ multiple copies of such elements.

In this contribution we set out to identify homologs of known exoribonuclease stalling elements and novel conserved structures. To this end, we computationally characterized homologs of experimentally verified xrRNA in tick-borne and no-known vector viruses that seem to form a coherent class of RNA structures with capability to stall exoribonucleases (T.xrRNA1/2, N.xrRNA). Likewise, we identified another class of xrRNAs in classic insect-specific flaviviruses (I.xrRNA1/2) which appears to be only distantly related to the former class. In the same line, we predicted a set of novel conserved elements in cISFVs that appear in quadruples and do not coincide with other insect-specific elements (I.Ra,I.Rb).

While we did not focus on studying the evolutionary history of these groups of elements in detail, our data suggests that many elements share ancestral roots. This is supported by the observation that at least the tick-borne, no-known-vector and *Aedes* spp. associated xrRNAs fold into a similar Y-shaped substructure, although the exact fold varies significantly among individual species.

We compiled a set of covariance models that can be used for rapid screening assays in the identification and characterization of novel flaviviruses. All models are available from GitHub via https://github.com/mtw/ITNFV-Data.

A major problem is the limited availability of diverse 3’UTR sequence data for many viruses analyzed here, particularly within cISFVs. Many novel ISFVs have previously been discovered, but 3’UTR sequences were only available for a subset of them. Future studies are required to shed more light on the evolutionary history of 3’UTR evolution in flaviviruses.

## Author Contributions

MTW conceived the study. RO and MTW analyzed the data, characterized conserved RNA elements and performed phylogenetic studies. All authors contributed to writing the paper and approved of the submitted version.

---

**Table A1.**
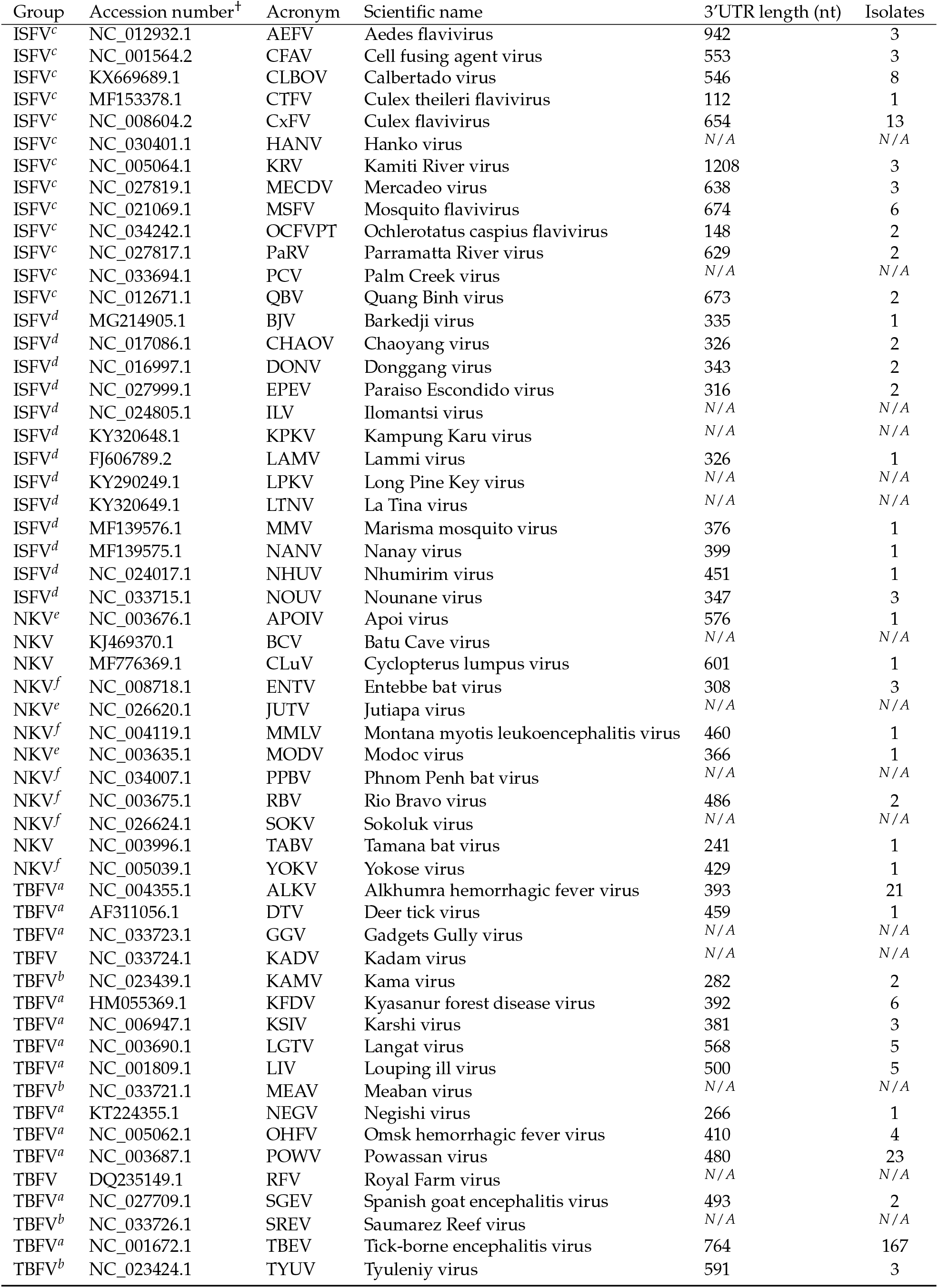
Viral genomes considered in this study. Flaviviruses are categorized into the groups tick-borne flaviviruses (TBFV), insect-specific flaviviruses (ISFV) and no-known-vector flaviviruses (NKV). The length of the 3’UTR is listed for each isolate. ^†^Representative accession number from the `refseq` database. Whenever a `refseq` genome was not available, the isolate with the longest 3’UTR was selected as representative species. ^*a*^Mammalian TBFVs. ^*b*^Seabird TBFVs. ^*c*^Classic ISFVs. ^*d*^Dual-host affiliated ISFVs. ^*e*^ Rodent-associated NKVs. ^*f*^ Bat-associated NKVs. ^*N/A*^ 3’UTR partial or not available in the refseq data set.

| Group | Accession number<sup>†</sup> | Acronym | Scientific name | 3'UTR length (nt) | Isolates |
| --- | --- | --- | --- | --- | --- |
| ISFV<sup>c</sup> | NC_012932.1 | AEFV | Aedes flavivirus | 942 | 3 |
| ISFV<sup>c</sup> | NC_001564.2 | CFAV | Cell fusing agent virus | 553 | 3 |
| ISFV<sup>c</sup> | KX669689.1 | CLBOV | Calbertado virus | 546 | 8 |
| ISFV<sup>c</sup> | MF153378.1 | CTFV | Culex theileri flavivirus | 112 | 1 |
| ISFV<sup>c</sup> | NC_008604.2 | CxFV | Culex flavivirus | 654 | 13 |
| ISFV<sup>c</sup> | NC_030401.1 | HANV | Hanko virus | N/A | N/A |
| ISFV<sup>c</sup> | NC_005064.1 | KRV | Kamiti River virus | 1208 | 3 |
| ISFV<sup>c</sup> | NC_027819.1 | MECDV | Mercadeo virus | 638 | 3 |
| ISFV<sup>c</sup> | NC_021069.1 | MSFV | Mosquito flavivirus | 674 | 6 |
| ISFV<sup>c</sup> | NC_034242.1 | OCFVPT | Ochlerotatus caspius flavivirus | 148 | 2 |
| ISFV<sup>c</sup> | NC_027817.1 | PaRV | Parramatta River virus | 629 | 2 |
| ISFV<sup>c</sup> | NC_033694.1 | PCV | Palm Creek virus | N/A | N/A |
| ISFV<sup>c</sup> | NC_012671.1 | QBV | Quang Binh virus | 673 | 2 |
| ISFV<sup>d</sup> | MG214905.1 | BJV | Barkedji virus | 335 | 1 |
| ISFV<sup>d</sup> | NC_017086.1 | CHAOV | Chaoyang virus | 326 | 2 |
| ISFV<sup>d</sup> | NC_016997.1 | DONV | Donggang virus | 343 | 2 |
| ISFV<sup>d</sup> | NC_027999.1 | EPEV | Paraíso Escondido virus | 316 | 2 |
| ISFV<sup>d</sup> | NC_024805.1 | ILV | Ilomantsi virus | N/A | N/A |
| ISFV<sup>d</sup> | KY320648.1 | KPKV | Kampung Karu virus | N/A | N/A |
| ISFV<sup>d</sup> | FJ606789.2 | LAMV | Lammi virus | 326 | 1 |
| ISFV<sup>d</sup> | KY290249.1 | LPKV | Long Pine Key virus | N/A | N/A |
| ISFV<sup>d</sup> | KY320649.1 | LTNV | La Tina virus | N/A | N/A |
| ISFV<sup>d</sup> | MF139576.1 | MMV | Marisma mosquito virus | 376 | 1 |
| ISFV<sup>d</sup> | MF139575.1 | NANV | Nanay virus | 399 | 1 |
| ISFV<sup>d</sup> | NC_024017.1 | NHUV | Nhumirim virus | 451 | 1 |
| ISFV<sup>d</sup> | NC_033715.1 | NOUV | Nounane virus | 347 | 3 |
| NKV<sup>e</sup> | NC_003676.1 | APOIV | Apoi virus | 576 | 1 |
| NKV | KJ469370.1 | BCV | Batu Cave virus | N/A | N/A |
| NKV | MF776369.1 | CLuV | Cyclopterus lumpus virus | 601 | 1 |
| NKV<sup>f</sup> | NC_008718.1 | ENTV | Entebbe bat virus | 308 | 3 |
| NKV<sup>e</sup> | NC_026620.1 | JUTV | Jutiapa virus | N/A | N/A |
| NKV<sup>f</sup> | NC_004119.1 | MMLV | Montana myotis leukoencephalitis virus | 460 | 1 |
| NKV<sup>e</sup> | NC_003635.1 | MODV | Modoc virus | 366 | 1 |
| NKV<sup>f</sup> | NC_034007.1 | PPBV | Phnom Penh bat virus | N/A | N/A |
| NKV<sup>f</sup> | NC_003675.1 | RBV | Rio Bravo virus | 486 | 2 |
| NKV<sup>f</sup> | NC_026624.1 | SOKV | Sokoluk virus | N/A | N/A |
| NKV | NC_003996.1 | TABV | Tamana bat virus | 241 | 1 |
| NKV<sup>f</sup> | NC_005039.1 | YOKV | Yokose virus | 429 | 1 |
| TBFV<sup>a</sup> | NC_004355.1 | ALKV | Alkhumra hemorrhagic fever virus | 393 | 21 |
| TBFV<sup>a</sup> | AF311056.1 | DTV | Deer tick virus | 459 | 1 |
| TBFV<sup>a</sup> | NC_033723.1 | GGV | Gadgets Gully virus | N/A | N/A |
| TBFV | NC_033724.1 | KADV | Kadam virus | N/A | N/A |
| TBFV<sup>b</sup> | NC_023439.1 | KAMV | Kama virus | 282 | 2 |
| TBFV<sup>a</sup> | HM055369.1 | KFDV | Kyasanur forest disease virus | 392 | 6 |
| TBFV<sup>a</sup> | NC_006947.1 | KSIV | Karshi virus | 381 | 3 |
| TBFV<sup>a</sup> | NC_003690.1 | LGTV | Langat virus | 568 | 5 |
| TBFV<sup>a</sup> | NC_001809.1 | LIV | Louping ill virus | 500 | 5 |
| TBFV<sup>b</sup> | NC_033721.1 | MEAV | Meaban virus | N/A | N/A |
| TBFV<sup>a</sup> | KT224355.1 | NEGV | Negishi virus | 266 | 1 |
| TBFV<sup>a</sup> | NC_005062.1 | OHFV | Omsk hemorrhagic fever virus | 410 | 4 |
| TBFV<sup>a</sup> | NC_003687.1 | POWV | Powassan virus | 480 | 23 |
| TBFV | DQ235149.1 | RFV | Royal Farm virus | N/A | N/A |
| TBFV<sup>a</sup> | NC_027709.1 | SGEV | Spanish goat encephalitis virus | 493 | 2 |
| TBFV<sup>b</sup> | NC_033726.1 | SREV | Saumarez Reef virus | N/A | N/A |
| TBFV<sup>a</sup> | NC_001672.1 | TBEV | Tick-borne encephalitis virus | 764 | 167 |
| TBFV<sup>b</sup> | NC_023424.1 | TYUV | Tyuleny virus | 591 | 3 |

